# GnotoBag: A versatile peat-based gnotobiotic system for plant microbiome research beyond model species

**DOI:** 10.64898/2026.11.22.768464

**Authors:** Nan Wu, Yu Ti Cheng, Bradley C. Paasch, Sheng Yang He

## Abstract

Plants and their microbial communities form a symbiotic system, where microbial communities play a key role in plant growth, development and stress resilience. However, the molecular mechanisms underlying plant-microbiota interactions remain poorly understood. A practical challenge is maintaining axenic conditions during prolonged plant growth while providing adequate space, ventilation, water, and nutrients. To advance research on plant-microbiome interactions, we developed the “GnotoBag” system, a bag-based extension of a previously established peat-based gnotobiotic plant growth platform. Key improvements of this GnotoBag system include enhanced ventilation and an integrated nutrient and water delivery system that sustains plant growth. These modifications support prolonged cultivation beyond the previously described 6.5-week workflow, enabling studies of microbiome effects on *Arabidopsis thaliana* growth and development under controlled microbial conditions, including the transition from vegetative to reproductive growth. Additionally, the expanded headspace of the GnotoBag provides sufficient space for cultivating larger plants, including rice and tomato, for at least four weeks. The increased headspace and the ability to add nutrients and water support research on larger crop plants, laying a foundation for future studies of microbiome functions in agricultural production.

## INTRODUCTION

Plants live in close association with diverse microbial communities that influence nutrient acquisition, growth, development, stress tolerance, and overall fitness [1–3]. To dissect these interactions mechanistically, plant researchers rely on gnotobiotic systems that allow plants to be cultivated under axenic conditions or with defined microbial strains or synthetic communities. Such systems are central to the study of plant-microbe and plant-microbiome interactions because they provide experimental reproducibility and precise control over microbial exposure.

A wide range of gnotobiotic plant growth systems has been developed, including agar-based platforms [4], hydroponic systems [5], fabricated microcosms [6], and substrate-based systems using inert or soil-like materials [4, 7, 8]. These approaches have enabled important advances in the study of microbial colonization, community assembly, host immune responses, and nutrient-related plant-microbiota interactions [5, 9–12]. However, many of these gnotobiotic plant growth systems were developed primarily for seedlings or short-term vegetative growth. As a result, plant growth can become constrained over time, and experimental conditions may diverge from those typically used in standard growth chambers or greenhouses. A well-designed gnotobiotic system must therefore balance microbial containment with plant growth requirements, including adequate space for root and shoot growth, gas exchange, temperature and humidity control, and sterile delivery of water or nutrients as required. Meeting these requirements becomes particularly important when studying biological processes that extend beyond the seedling stage. Developmental transitions such as bolting, flowering, reproductive growth, and seed production require longer cultivation periods than are supported by many commonly used gnotobiotic systems. Similar challenges arise when cultivating crop species such as rice and tomato, which often require greater growth space and longer cultivation periods than *Arabidopsis*. Thus, although existing systems are highly effective for many short-term applications, there remains a need for a gnotobiotic platform that supports prolonged plant growth and later developmental stages without compromising microbial control.

To address this need, we developed GnotoBag, a modular bag-based gnotobiotic cultivation system adapted from an earlier peat-based platform [8]. GnotoBag is designed to support longer-term plant growth through improved ventilation, expanded headspace, and sterile delivery of water and nutrients, while providing flexibility to accommodate different substrates, pot sizes, plant species, and experimental designs (Fig. 2a–d). In this protocol, we describe the design, assembly, sterilization, planting, irrigation, and maintenance of the GnotoBag system. We demonstrate its utility for controlled microbiome studies across different developmental stages and experimental timescales using *Arabidopsis*, rice, and tomato (Figs. 3–6). We also discuss practical considerations for maintaining sterility during extended cultivation.

**Figure 1:**
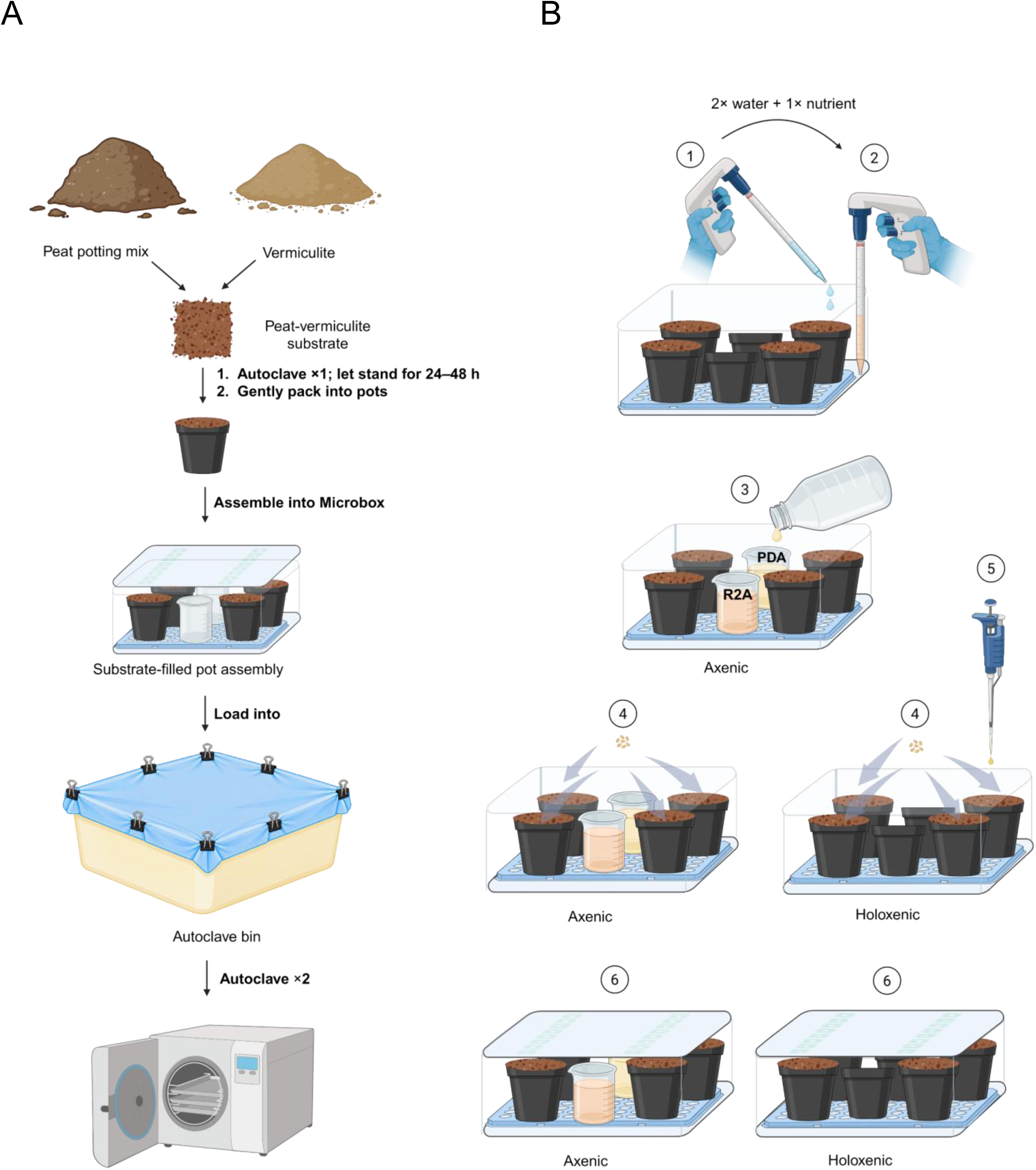
Schematic illustration of the substrate-filled pot workflow for Microbox-based gnotobiotic cultivation. (a) Preparation of substrate-filled pots. A loose peat–vermiculite substrate is autoclaved to reduce microbial load and gently packed into 2-inch polypropylene pots, and the packed pots are assembled in Microboxes. The assembled units are then sterilized by two autoclave cycles spaced 24 to 48 h apart. (b) Post-sterilization preparation, sowing, and microbial inoculation workflow. After the second autoclave cycle, the assembled units are transferred to a biosafety cabinet or laminar flow hood. In Steps 1 and 2, the sterilized substrate-filled pots are flushed with warm, autoclaved water twice, followed by warm, autoclaved nutrient solution once. In Step 3, Reasoner’s 2A (R2A) agar and potato dextrose agar (PDA) media are dispensed into sterility-monitoring beakers in boxes designated for axenic conditions while the system cools to room temperature. In Step 4, surface-sterilized seeds are sown onto the substrate surface in each pot. In Step 5, each pot receives 1 mL of the prepared microbial inoculum or the corresponding axenic control preparation. In Step 6, the Microbox lids are closed with the support racks remaining beneath the pots, and the units are transferred to a growth chamber with the desired light and temperature settings.

**Figure 2:**
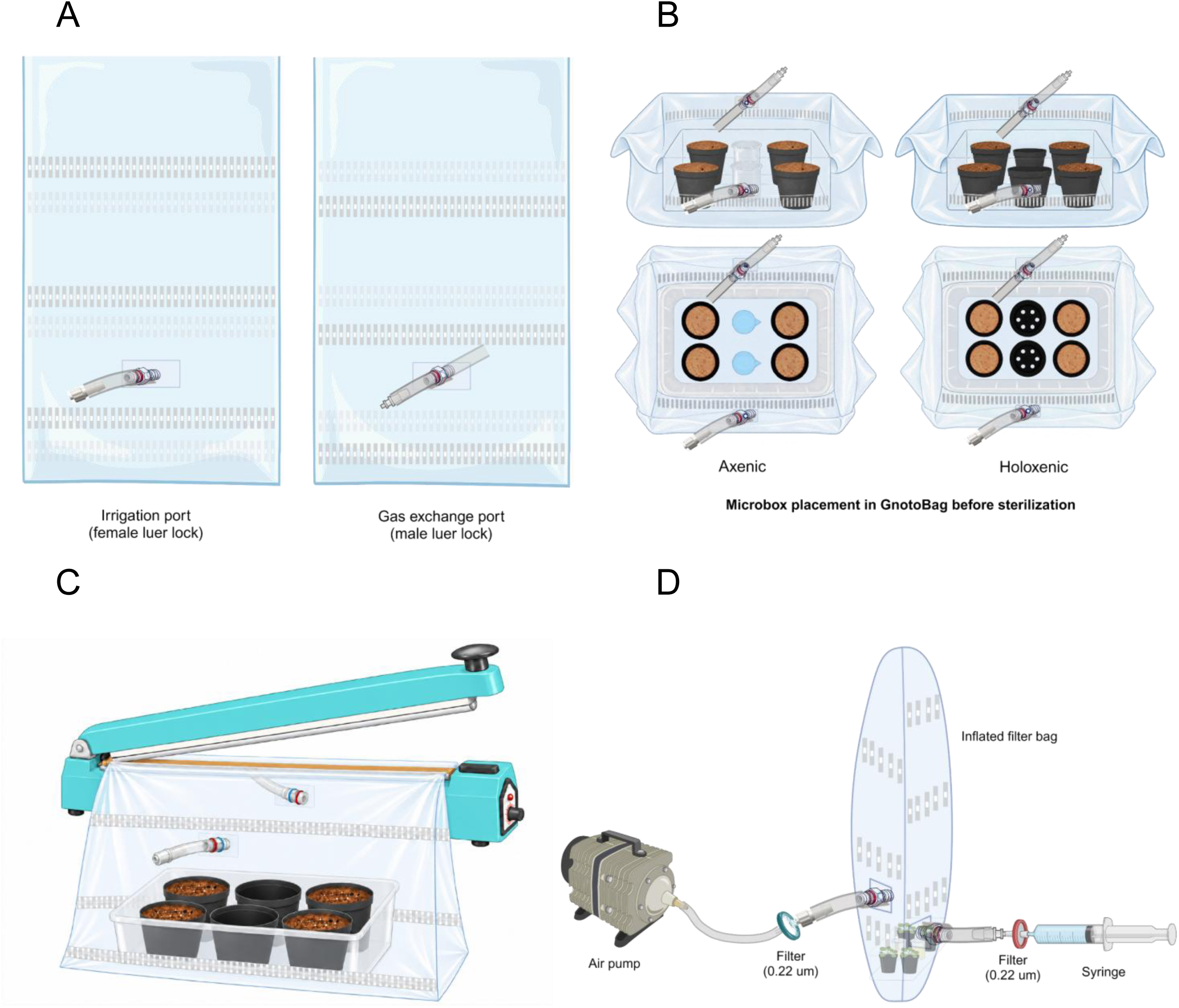
Schematic overview of the GnotoBag assembly and operation workflow. (a) Preparation of the polypropylene (PP) zipper filter bag. One reinforced port is punched on each side of the bag, and the ports are fitted with the corresponding barbed and luer-lock connectors. (b) Placement of the Microbox assembly into the PP zipper filter bag before sterilization. The Microbox containing substrate-filled pots is positioned at the center of the bag bottom. To reduce the overall height of the assembly before autoclaving, the upper portion of the PP zipper filter bag is gently rolled downward around the outside of the Microbox until the folded edge is approximately level with the top rim of the box. The assembly is then processed through the substrate-filled pot preparation and sterilization workflow shown in Fig. 1. (c) Heat sealing of the expanded GnotoBag. After the substrate-filled pot preparation, sterilization, flushing, sowing, and inoculation steps are completed, the previously rolled-down upper portion of the bag is carefully unfolded from around the Microbox and extended upward to restore the bag to its fully expanded configuration. The open upper end of the bag is positioned between the jaws of a heat sealer, with the bag film kept flat and evenly aligned. The bag is then heat-sealed to close the opening and generate a continuous, gap-free seal. (d) Final setup and operation of the GnotoBag system. Inside a biosafety cabinet or laminar flow hood, sterile 0.22 µm filters are attached to both luer ports. The assembled GnotoBag is then transferred to a growth chamber or greenhouse. One port is connected to an air pump through aquarium airline tubing for continuous aeration, whereas the opposite port can be connected to a syringe for sterile water or nutrient supplementation. The GnotoBag typically inflates within approximately 10 min, after which continuous aeration is maintained throughout plant growth.

**Figure 3:**
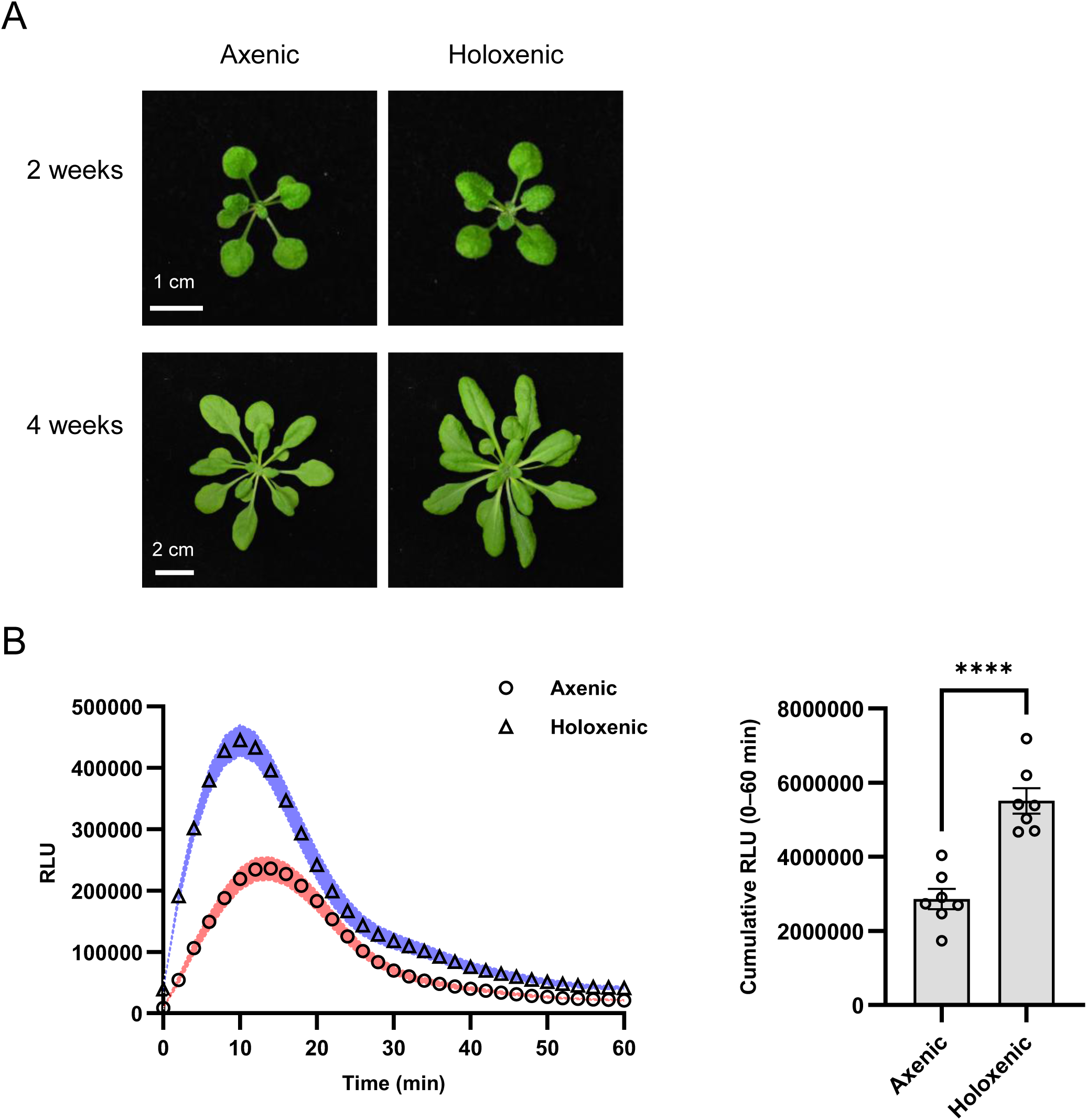
*A. thaliana* growth and flg22-induced ROS burst in substrate-filled pots maintained within Microboxes. (a) Representative images of *A. thaliana* plants grown in substrate-filled pots under axenic or holoxenic conditions at 2 and 4 weeks after sowing. Holoxenic plants were inoculated with 1 mL of SynCom^Col-0^ prepared as described in this protocol. Axenic controls received 1 mL of the corresponding heat-killed SynCom^Col-0^. (b) flg22-induced ROS burst in 4-week-old Col-0 plants grown under axenic or holoxenic conditions. Leaf discs were treated with 100 nM flg22, and ROS production was monitored over 60 min as relative luminescence units (RLU) (left). Shaded bands indicate SEM. Total ROS production was calculated over the 60-min assay period (right). Points overlaid on the bar plot indicate individual biological replicates (n = 7 per condition); bars and error bars indicate mean ± SEM. Statistical significance for total ROS production was determined using a two-tailed unpaired t-test with Welch’s correction. Asterisks indicate a significant difference between axenic and holoxenic plants; ****, P < 0.0001.

**Figure 4:**
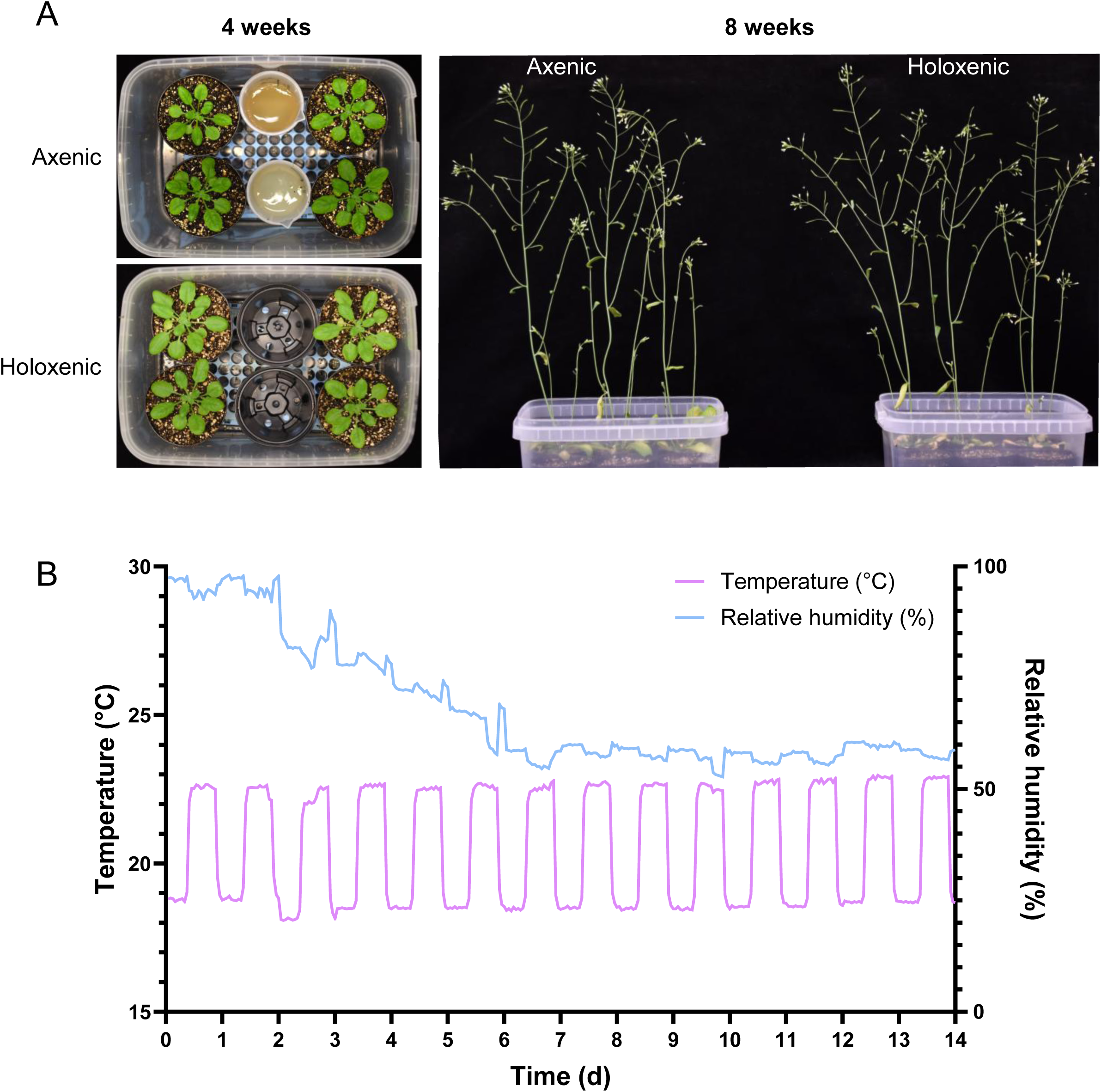
*A. thaliana* growth and environmental monitoring in the GnotoBag system under axenic or holoxenic conditions. (a) Representative images of *A. thaliana* plants grown in the GnotoBag system under axenic or holoxenic conditions at 4 weeks after sowing (left) and 8 weeks after sowing (right). Plants were grown inside sealed GnotoBags. The Microboxes containing the plants were removed from the bags for imaging. Holoxenic plants were inoculated with 1 mL of SynCom^Col-0^ prepared as described in this protocol. Axenic controls received 1 mL of the corresponding heat-killed SynCom^Col-0^. (b) Temperature and relative humidity inside the GnotoBag system during plant growth. Temperature and relative humidity were continuously monitored using a sensor placed inside the GnotoBag during the first 2 weeks after system establishment. The growth chamber setpoints were 20 °C and 40% relative humidity during the light period and 18 °C and 40% relative humidity during the dark period.

**Figure 5:**
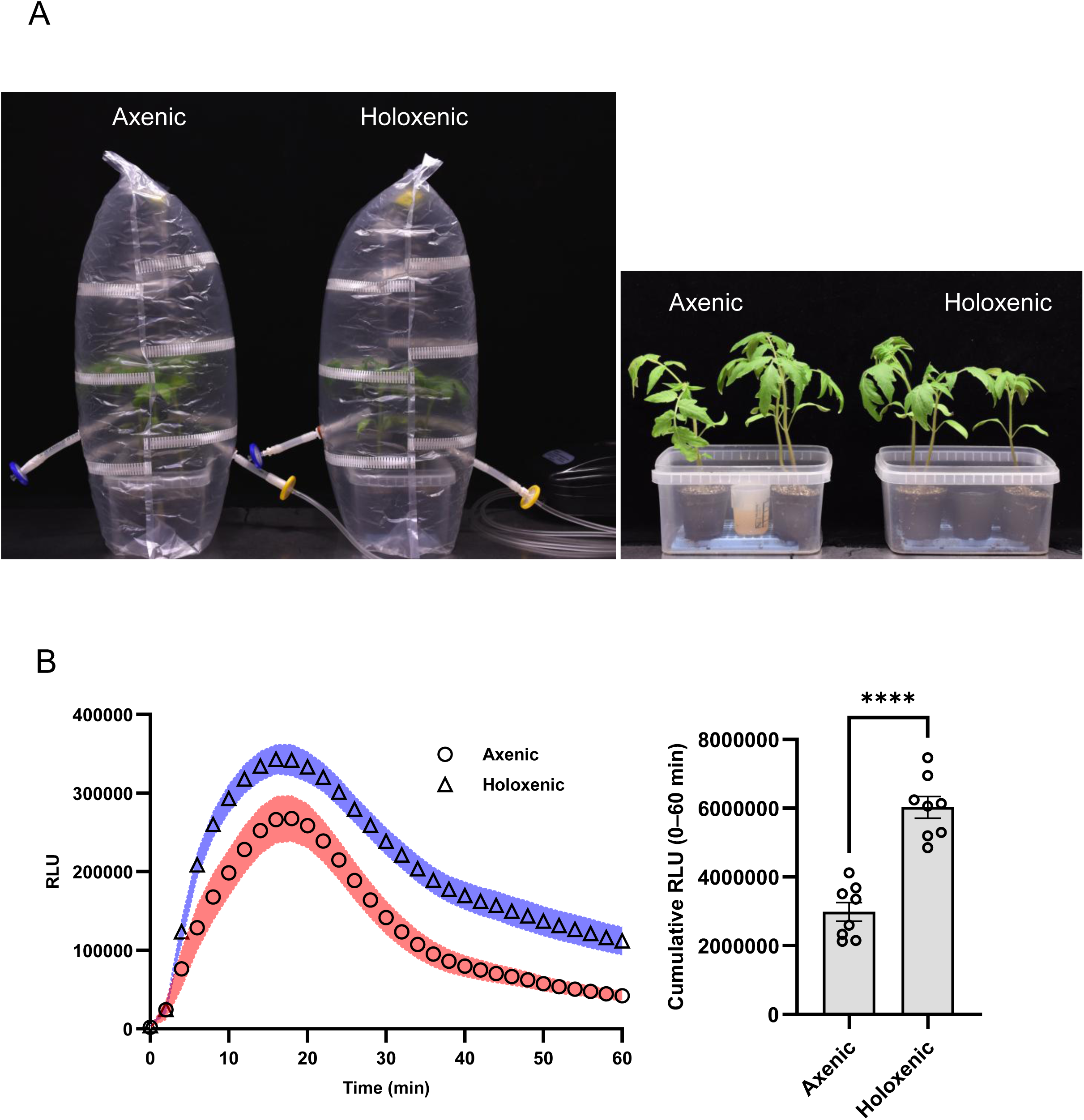
Tomato growth and flg22-induced ROS burst in the GnotoBag system. (a) Tomato cultivar Moneymaker growth and available headspace within the GnotoBag system. Representative 4-week-old plants are shown inside the GnotoBag system to illustrate the available headspace for continued growth (left). The same plants are shown after removal from the system to show overall plant size and morphology (right). Holoxenic plants were inoculated with 1 mL of soil slurry prepared as described in this protocol. Axenic controls received 1 mL of the corresponding heat-killed soil slurry. (b) flg22-induced ROS burst in 4-week-old tomato plants grown under axenic and holoxenic conditions. Leaf discs were treated with 100 nM flg22, and ROS production was monitored over 60 min as RLU (left). Shaded bands indicate SEM. Total ROS production was calculated over the 60-min assay period (right). Points overlaid on the bar plot indicate individual biological replicates (n = 8 per condition). Bars and error bars indicate mean ± SEM. ****, P < 0.0001.

**Figure 6:**
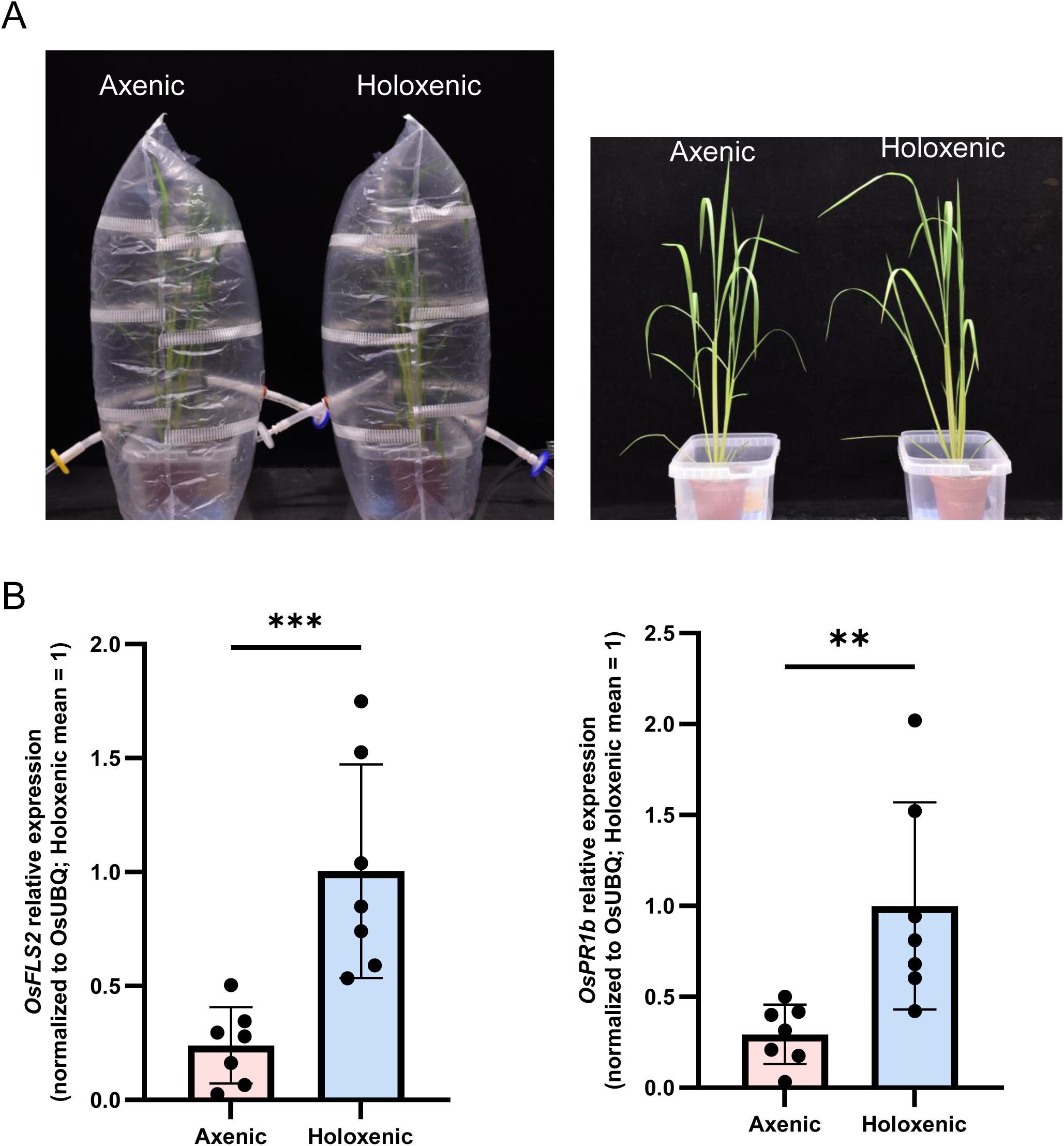
Rice plants grown in the GnotoBag system under axenic or holoxenic conditions. (a) Rice cultivar Kitaake growth and available headspace within the GnotoBag system. Representative 4-week-old rice plants are shown inside the GnotoBag system to illustrate the available headspace for continued growth (left). The same plants are shown after removal from the system to show overall plant size and morphology (right). For holoxenic conditions, each pot was inoculated with 1 mL of soil slurry prepared as described in this protocol. Axenic controls received 1 mL of the corresponding heat-killed soil slurry. (b) RT-qPCR analysis of defense-associated marker gene expression in rice leaves from plants grown in the GnotoBag system under axenic and holoxenic conditions. Transcript levels of *OsFLS2* and *OsPR1b* were normalized to *OsUBQ*. For each gene, expression values were scaled to the mean expression level of the holoxenic condition, which was set to 1. Each dot represents an individual biological replicate (n = 7 per condition). Bars show mean ± SD. Statistical significance was determined using a two-tailed Mann–Whitney U test. **, P < 0.01, ***, P < 0.001.

## EQUIPMENT AND MATERIALS

### Equipment

A biosafety cabinet or a laminar flow hood

Controlled environment chamber with desired lighting (e.g., Percival, cat. no. CU36L5 or BioChambers, FXC-19-GD)

Tabletop impulse sealer (12- to 16-inch sealing length, e.g., AIE-300 or AIE 400P)

Electronic pipette controller (Pipet-Aid XP, Drummond Scientific Company)

Analytical balance (Mettler Toledo, model AB54)

Gravity-displacement steam autoclave (AMSCO, Eagle Series, model 3021)

Wireless temperature and relative humidity sensor (SensorPush, model HT1)

### Materials

Substrate components: peat potting mix (e.g., Sun Gro Horticulture, Sunshine Mix #5) and medium-grade vermiculite (P.V.P. Industries, Inc., North Bloomfield, OH, USA)

Nalgene large polypropylene sterilizing pans (20 L, Thermo Scientific, cat. no. 6900-0020) 2-inch polypropylene nursery pots (Amazon, ASIN B00LH1NMV0)

Microbox container, #40 green filter, autoclavable (SacO2, cat. no. TP1600+TPD1600 #40 (NG/NP))

Polypropylene Zipper Microsac (SacO2, cat. no. PPD50/SEH6/V32-49) 50-mL polypropylene beakers (Thermo Scientific, cat. no. 1201-0050)

50-mL individually wrapped sterile serological pipette (CELLTREAT Scientific, cat. no. 229230B)

Cell strainer, 70 µm (CELLTREAT Scientific, cat. no. 229483)

Cell strainer, 40 µm (CELLTREAT Scientific, cat. no. 229481)

Sterilization wraps 30-inch × 30-inch (Medline, cat. no. GEM1130S) or 24-inch × 24-inch (Medline, cat. no. GEM1124S)

Pipettes with sterile 1-mL filtered and 250-µL wide-orifice tips (e.g., Rainin, cat. no. 30389212 and 30389249)

Air pumps (110 GPH 40-watt 8-outlet commercial air pump for aquarium and hydroponic systems)

Tubing (Cole-Parmer, Thermoplastic (SEBS) Tubing, 1/4-inch ID × 3/8-inch OD; 50 ft; Penn-Plax Standard)

Airline tubing for aquariums – clear and flexible (Amazon)

Sterile hydrophilic syringe filters for irrigation (0.22 µm, 33 mm diameter, mixed cellulose esters [MCE] membrane; Millex-GS, MilliporeSigma, cat. no. SLGS033SS)

Sterile syringe filters used on the gas-exchange line (0.22 µm, 33 mm diameter, hydrophilic polyvinylidene fluoride [PVDF] membrane; Millex-GV, MilliporeSigma, cat. no. SLGVR33RB)

3M Scotch heavy duty packaging tape

60-mL luer-lock syringes

Male luer (1/4-inch ID barb, straight nylon fitting, Value Plastics, part no. MTLL055-1)

Female luer (1/4-inch ID barb, straight polypropylene fitting, Value Plastics, part no. FTLL055-6005)

Barbed bulkhead fitting (polypropylene, for 6.5-mm ID tubing, Thermo Scientific Nalgene, cat. no. 6149-0002)

Autoclavable plastic spacer/support rack (“blue rack”) – polypropylene or polycarbonate insert from pipette tip boxes (USA Scientific, part of the Pipette Tip Refills, cat. no. 1111-2721)

Handheld round hole punch (1/4-inch diameter)

### Reagents

Sterile Milli-Q water (reverse osmosis filtered, or an equivalent quality water)

Linsmaier and Skoog (LS) basal salts (pH-buffered to 5.7 ± 0.2, Caisson Labs, cat. no. LSP03)

Reasoner’s 2A (R2A) agar (MilliporeSigma, cat. no. 17209)

Potato dextrose agar (PDA; MilliporeSigma, cat. no. 70139)

Bleach (common household bleach, 5.25–8.25% sodium hypochlorite, e.g., Clorox)

0.1% (w/v) agar (autoclaved, BioBasic, CAS no. 9002-18-0)

75% (v/v) ethanol for disinfection

### Procedure 1: Substrate adaptation and preparation of substrate-filled pots (Fig. 1)

In this procedure, we describe the adaptation and preparation of a loose peat-vermiculite substrate for gnotobiotic plant growth. In the original GnotoPot workflow, compressed Jiffy-7 peat pellets were used as the growth substrate. During the development of the GnotoBag workflow, we found that the fixed size and shape of these pellets limited their use with different pot sizes, seed types, and larger plant species. We also encountered variability among pellet lots. We therefore adopted a loose peat-based potting mix supplemented with vermiculite, which can be packed into polypropylene nursery pots of different sizes and assembled within Microboxes for subsequent gnotobiotic cultivation (Fig. 1a).

### Seed preparation (Total elapsed time 3 days; hands-on time ≤ 1 h)

- Prepare surface-sterilized *Arabidopsis* seeds as described by Kremer et al. [8].
- Prepare surface-sterilized tomato seeds with modifications from a previously described protocol [13]. Briefly, pre-soak dry tomato seeds overnight in sterile distilled water to allow full imbibition. Treat the imbibed seeds with 1% sodium hypochlorite solution for 15 min with gentle mixing. After sterilization, rinse the seeds five times with sterile distilled water under aseptic conditions to remove residual bleach, and use them immediately for sowing.
- Prepare surface-sterilized rice seeds with modifications from the Kitaake rice transformation protocol developed by the Langdale Lab (https://langdalelab.com/protocols/transformation/).

Sterilization of the substrate and assembly of substrate-filled pots (Total elapsed time ∼3–6 days) (Fig. 1a)

1. The following quantities are sufficient for preparing approximately thirty-two 2-inch substrate-filled pots. Thoroughly blend 1.5 L of loosely packed peat-based potting mix with 1.5 L of medium-grade vermiculite. Moisten the mixture evenly with 1 L of distilled water. Distribute the substrate on clean large Nalgene polypropylene sterilizing pans at a depth of approximately 2 inches. Cover each pan with one layer of sterilization wrap and secure the wrap with binder clips. Autoclave for 45 min on liquid cycle (121 °C, 15 psi, slow exhaust with forced liquid cooling) and immediately remove the pans from the autoclave after the cycle is complete. Let sit covered with sterilization wrap at room temperature (RT; 22–25 °C) for 24–48 h (Hands-on time ∼60 min).

Note: Do not let materials sit in the autoclave machine after cycling as this may lead to excessive drying or structural disintegration of the substrate that results in suboptimal plant growth.

2. Assembly of substrate-filled pots (Hands-on time ∼1 h). Mix the sterilized substrate thoroughly to achieve a consistent composition. Fill each pot to the rim with the prepared peat-based substrate. Gently tap the pot 1 to 2 times on a firm surface to eliminate large air pockets and allow the substrate to settle evenly (do not compress the substrate manually). After tapping, ensure that the substrate remains level with the rim. Weigh each filled pot and adjust the amount of substrate by adding or removing material as needed to reach the predetermined target net substrate weight for that pot size, within ±1.5% of the target. This step helps ensure consistent substrate volume and packing density across pots.

Note: Based on our experience, filling one 2-inch pot with the prepared substrate described above requires approximately 30.0 ± 0.45 g of substrate. All subsequent volume calculations described below are based on the use of 2-inch pots.

3. Place one autoclave-compatible plastic spacer/support rack (“blue rack”) at the bottom of each Microbox to elevate the pots, allow excess liquid to drain beneath them, and prevent waterlogging. Place four substrate-filled pots in the outer positions of each Microbox, leaving the two central positions available for sterility-monitoring beakers or empty pots. For axenic conditions, place two empty beakers in the central positions for subsequent addition of R2A and PDA agar to monitor bacterial and fungal contamination, respectively (Fig. 1b, step 3). For holoxenic conditions, replace the beakers with two empty pots. Place the lid loosely on each Microbox without snapping it closed around the entire rim. Place the Microboxes containing substrate-filled pots into clean Nalgene polypropylene sterilizing pans, cover each sterilizing pan with three layers of sterilization wrap, and secure the wraps with binder clips.

Note: Do not snap the Microbox lid closed around the entire rim before autoclaving, as pressure changes during autoclaving may deform the containers and lids. Additional information on autoclaving Microboxes is available on the manufacturer’s website.

4. Autoclave the assembled units for 45 min on liquid cycle (121 °C, 15 psi, slow exhaust with forced liquid cooling) and unload the assembled units immediately after the autoclave cycle is complete. Let sit covered with sterilization wrap at RT for 24–48 h.
5. Autoclave the assembled units for a second time (to further reduce the risk of spore contamination) for 45 min on liquid cycle (121 °C, 15 psi, slow exhaust with forced liquid cooling).

### Preparation of flushing and irrigation solutions and sterility-monitoring media (Hands-on time ∼30 min)

6. For flushing and irrigation, prepare the required volumes of Milli-Q water and 0.5× LS nutrient solution. Use two volumes of Milli-Q water and one volume of 0.5× LS nutrient solution for each pot. For example, one pot containing 30.0 g of net substrate requires approximately 60 mL of Milli-Q water and 30 mL of 0.5× LS nutrient solution. Autoclave the solutions for 45 min. After autoclaving, transfer them to a pre-cleaned laminar flow hood or biosafety cabinet.
7. For axenic conditions, prepare R2A and PDA media to monitor sterility. Prepare approximately 30 mL of each medium per unit. Autoclave the media as required. After autoclaving, keep the R2A and PDA media molten in a 65 °C water bath until use.

Flushing, inoculation, and sowing of substrate-filled pots (Hands-on time ∼3 h for 8 Microboxes) (Fig. 1b)

Note: Clean the work surface of a laminar flow hood or biosafety cabinet before use. Use aseptic technique throughout the procedure to prevent microbial contamination.

8. After the second autoclave cycle, place the autoclaved pans containing Microboxes with substrate-filled pots into a pre-cleaned laminar flow hood or biosafety cabinet. In the hood, carefully transfer the loosely covered Microboxes out of the pans.
9. While the substrate-filled pots are still warm, carefully open the Microbox lids. Using a sterile serological pipette, gently apply one volume of warm (∼60 °C), autoclaved Milli-Q water to the surface of each pot. After all pots in the Microbox have been flushed, remove the flow-through liquid that has accumulated at the bottom of the Microbox using a sterile serological pipette. Repeat this flushing procedure once, for a total of two washes per pot. After the two Milli-Q water flushes, irrigate each pot once with one volume of warm, autoclaved 0.5× LS nutrient solution. Remove any excess flow-through liquid from the bottom of the Microbox if necessary (Fig. 1b, steps 1 and 2).

Note: Dispense water and nutrient solution slowly and evenly across the substrate surface to allow the liquid to percolate through the substrate. This ensures thorough rehydration and helps flush out potential phytotoxic byproducts generated during autoclaving. Avoid rapid dispensing to prevent overflow or channeling.

10. After completing the flushing and irrigation steps, and while the Microboxes are cooling to room temperature, dispense ∼30 mL of molten R2A medium into one sterility-monitoring beaker and ∼30 mL of molten PDA medium into the other (Fig. 1b, step 3). Allow the media to solidify and the substrate-filled pots to cool to room temperature before sowing seeds and applying microbial inoculum.
11. For *Arabidopsis* grown in Microboxes, sow 3 to 5 surface-sterilized seeds onto the substrate surface in each pot using sterile 1-mL tips or 250-µL wide-orifice tips (Fig. 1b, step 4). Apply 1 mL of the prepared microbial community inoculum to the substrate surface in each pot for holoxenic conditions (Box 1 and Fig. 1b, step 5), or 1 mL of the corresponding control preparation described in Box 1 for axenic conditions. For both conditions, snap each Microbox lid closed around the entire rim (Fig. 1b, step 6). Transfer the closed Microboxes from the laminar flow hood or biosafety cabinet to the plant growth chambers.

### Plant care and sterility testing during Microbox growth

11. For axenic conditions, confirm sterility before using plant material for experiments (see Box 2).
12. Plants grown in Microboxes require minimal care aside from thinning, when applicable, and sterility testing before experiments. Regularly rotate Microboxes within the growth chamber to minimize positional effects on plant growth.

**Box 1 Preparation of microbial community for inoculation**

##### 1.1 Undefined natural microbiota: soil slurry

1. Add 5 g of soil to 100 mL of sterile Milli-Q water in a clean container. Stir the mixture at RT for 20 min at 200–300 rpm using a magnetic stir plate. After stirring, to remove large particulates, sequentially filter the soil slurry through a 70 µm cell strainer followed by a 40 µm cell strainer into a clean media bottle.

Note: Divide the prepared soil slurry into two equal aliquots. Retain one aliquot without further treatment for holoxenic inoculation and autoclave the other at 121 °C for 45 min to prepare the heat-killed control. Allow the autoclaved aliquot to cool to room temperature.

2. For holoxenic conditions, apply 1 mL of the prepared natural microbial community inoculum to the substrate surface in each pot. For axenic controls, prepare a corresponding heat-killed soil slurry and apply 1 mL to the substrate surface in each pot.
3. Use the remaining inoculum to perform serial dilution for quantification. Plate appropriate dilutions on R2A agar to estimate the number of colony-forming units (CFUs) inoculated onto each pot.

##### 1.2 Defined microbiota: synthetic community (SynCom)

1. Inoculate each bacterial strain from frozen glycerol stock onto R2A agar plates and incubate at RT. Monitor bacterial growth periodically. Transfer plates with visible lawns to 4 °C to slow further growth, while allowing slower-growing strains to continue incubating at RT until sufficient biomass can be obtained.
2. Scrape bacterial lawns from R2A plates and resuspend each strain in approximately 2 mL of sterile 10 mM MgCl₂. Mix thoroughly to ensure an even suspension. Take an aliquot from each suspension to measure its optical density at 600 nm (OD₆₀₀); normalize each suspension with sterile 10 mM MgCl₂ to OD₆₀₀ = 0.4. Combine equal volumes of the normalized bacterial suspensions in a sterile 50-mL tube.
3. Add an equal volume of 10% dimethyl sulfoxide (DMSO) to the combined and normalized bacterial suspension to achieve a final suspension with OD₆₀₀ = 0.2 in 5% DMSO. Mix gently but thoroughly to ensure uniform distribution. Aliquot 250–300 µL of the prepared SynCom into sterile microcentrifuge tubes. Snap-freeze the aliquots in liquid nitrogen and store at −80 °C until use.
4. To use the prepared SynCom, remove an aliquot from the −80 °C freezer and thaw it in a 37 °C water bath with gentle agitation for 1 min. Dilute the suspension to the desired concentration using sterile 10 mM MgCl₂. For SynCom^Col-0^ [14], use a 100,000-fold dilution of the frozen stock prepared above. Divide the diluted suspension into two equal aliquots. Retain one aliquot without further treatment for holoxenic inoculation and autoclave the other at 121 °C for 45 min to prepare the heat-killed control. Allow the autoclaved aliquot to cool to room temperature. Under aseptic conditions, apply 1 mL of the live SynCom suspension or the corresponding heat-killed control to the substrate surface in each pot for holoxenic or axenic conditions, respectively.
5. Prepare serial dilutions of the remaining diluted SynCom inoculum for quantification. Plate appropriate dilutions on R2A agar to estimate the number of CFUs applied per pot for each inoculation.

**Box 2 Assessment of sterility in gnotobiotic systems** (Hands-on time ∼50 min for 8 Microboxes containing substrate-filled pots)

For plants grown in Microboxes without GnotoBags (Procedure 1), assess sterility at the time of thinning (∼10-day-old seedlings) or ∼7 days before the planned experiment.

Note: All steps described below should be performed in a pre-cleaned laminar flow hood or biosafety cabinet using aseptic techniques to minimize the risk of contamination.

1. In a pre-cleaned laminar flow hood or biosafety cabinet, open Microboxes containing substrate-filled pots. Thin seedlings using sterile tweezers and place the removed seedlings onto R2A and PDA agar plates to assess microbial contamination. Incubate the plates at RT for at least 1 week and monitor for microbial contamination.
2. Disinfect tweezers between Microboxes by immersing them in diluted bleach solution containing 0.5–1% sodium hypochlorite for 1 min, followed by two rinses with autoclaved Milli-Q water.
3. To assess sterility before planned experiments, open Microboxes containing substrate-filled pots in a pre-cleaned laminar flow hood or biosafety cabinet. Transfer 1 mL of standing liquid from the bottom of each Microbox onto R2A and PDA agar plates. Using a sterile inoculation loop, collect a small amount of substrate and spread it evenly onto separate R2A and PDA agar plates. Incubate the plates at RT for at least 1 week and monitor for microbial contamination.

Note: For GnotoBag units (Procedure 2), monitor the internal R2A and PDA agar-containing beakers for visible signs of contamination without opening the bags during cultivation. Collect samples for an endpoint sterility check at the final experimental harvest, as described in Procedure 2. Determine whether the corresponding plant samples and data can be included in the final analysis after at least 1 week of incubation.

### Procedure 2: GnotoBag system (Fig. 2)

Construction of the GnotoBag system (Total elapsed time ∼2 h for 12 bags) (Fig. 2a)

Note: GnotoBags can be reused up to three times. The bag assembly procedure only needs to be performed during the initial construction. Before reuse, inspect each bag and connector for visible damage, deformation, or leakage.

*Port-site definition and aperture creation*

1. *Prepare reinforced port sites*. Place a new polypropylene (PP) zipper filter bag flat on a clean bench surface, with the sealed bottom edge facing you and the zipper opening facing away from you. On one side of the bag, mark a position approximately 6 inches from the bottom edge, avoiding the filter strips. Apply a 5 × 5 cm piece of transparent heavy-duty Scotch tape to the inner surface of the bag at this position, and apply a second 5 × 5 cm piece of tape to the corresponding position on the outer surface. The two tape pieces should be aligned so that the bag film is sandwiched between them, forming a reinforced tape-film-tape patch. Repeat the same procedure at the corresponding position on the opposite face of the bag to generate a second reinforced patch. These reinforced patches provide mechanical support for subsequent hole punching, reduce tearing of the bag film, and produce a clean hole edge.
2. *Punch the ports*. Firmly press the handles of a 1/4-inch (6.35 mm) handheld hole punch to make a clean, circular hole in the center of each reinforced tape-film-tape patch.

*Bulkhead fitting installation and luer/tubing assembly*

3. *Insert barbed bulkhead fittings*. Insert one barbed bulkhead fitting through each reinforced hole from the outside of the bag toward the interior, with the wide flat flange positioned inside the bag. Ensure that the flange sits flush against the inner surface of the bag film.
4. *Attach external tubing and luer connectors*. On the exterior barb of each bulkhead fitting, push-fit a ∼5 cm segment of 1/4-inch ID SEBS tubing. Attach a luer-lock connector to the distal end of each tubing segment, with one face of the bag terminating in a female luer-lock connector and the opposite face of the bag terminating in a male luer-lock connector. On the internal barb of the irrigation port, attach a ∼5 cm segment of tubing to serve as a guide tube for directing irrigation solution onto the substrate.

Note: Specify the connector type for each port at this step to avoid ambiguity later. Designate one port as the <u>irrigation port</u> and terminate its external tubing with a **female luer-lock connector.** This port is used for introducing liquid into the bag via a sterile filter unit. Designate the other port as the <u>gas exchange</u> <u>port</u>, terminate its external tubing with a **male luer lock**, and use it for connection to the air pump through a sterile air filter.

### Sterilization and assembly of substrate-filled pots in GnotoBags (Hands-on time: ∼3–6 h) (Fig. 2b)

5. *Load substrate-filled pots into Microboxes.* Assemble Microboxes containing substrate-filled pots as described in Procedure 1.
6. *Bag the Microbox before autoclaving.* Place the entire Microbox into the prepared PP zipper filter bag. Loosely place a lid on top of the Microbox to reduce excessive drying during sterilization. Roll the open rim of the bag down to approximately the same height as the Microbox so that the bag loosely surrounds the unit without enclosing it. Do not stretch the bag tightly across the lid or block any vents or filter elements.
7. Place the assembled bagged units into large Nalgene polypropylene sterilizing pans, keeping them upright without stacking or tilting them. Because the Microboxes enclosed in GnotoBags are bulkier than unbagged Microbox assemblies, load no more than four bagged units per pan. Autoclave the loaded pans twice using the cycle described in Procedure 1. After the second cycle, carefully unload the covered pans and transfer them to a pre-cleaned laminar flow hood or biosafety cabinet.

### Irrigation, inoculation, and sowing in GnotoBags (Total elapsed time ∼3–6 h)

Note: All steps described below should be performed in a pre-cleaned laminar flow hood or biosafety cabinet using aseptic techniques to minimize the risk of contamination.

8. After autoclaving, perform flushing, initial irrigation, sterility-monitoring media preparation, seed sowing, and microbial inoculation as described in Procedure 1.

Note: When preparing GnotoBag units, sow only one seed per pot, as the bags remain sealed for the duration of plant growth and do not permit thinning. Use high-quality seed stocks to ensure reliable germination.

### Final placement, heat sealing, and sterile filter connection (Fig. 2c, d)

9. After flushing, seed sowing, and application of the microbial inoculum or corresponding axenic control preparation, ensure that the Microbox lid is removed and leave the Microbox uncovered inside the GnotoBag. Fully unfold the bag and position the Microbox at the center of the bag bottom.
10. Heat-seal the bag. Use a tabletop impulse heat sealer to seal the open end of the bag, ensuring a continuous, gap-free seal. Inspect the seal for microchannels or incomplete fusion, and reseal if necessary. If desired, add a second parallel seal slightly offset from the first.
11. Attach sterile 0.22 µm filters to both luer ports. For the air line, connect the filter’s female luer lock to the male luer port on the GnotoBag. For the irrigation line, connect the filter’s male luer lock to the female luer port on the GnotoBag. Confirm that both filters are securely attached.

Note: Install a hydrophilic 0.22 µm inline filter (suitable for aqueous media, e.g., MCE) on the irrigation line. Do not use hydrophobic membrane filters as they impede liquid flow.

12. Carefully transfer the sealed GnotoBag to the growth chamber. Connect the air pump to the gas-exchange port via aquarium airline tubing and a sterile 0.22 µm Millex-GV PVDF syringe filter (MilliporeSigma, cat. no. SLGVR33RB). Maintain continuous sterile airflow throughout plant growth.

### Plant care and sterility monitoring during gnotobiotic growth

13. Temperature and relative humidity within the GnotoBag change during cultivation and should be monitored, as conditions inside the bag may differ from the growth chamber settings (Fig. 4b). Under these conditions, plants continue to transpire and require regular watering during extended growth. To water plants in the GnotoBag system, temporarily disconnect the air-pump tubing from the upstream side of the air filter, leaving the filter securely attached to the GnotoBag gas-exchange port throughout irrigation. Use a 60-mL luer-lock syringe to draw up room-temperature autoclaved water. Connect the syringe to the hydrophilic 0.22 µm MCE filter on the irrigation line via the luer-lock fitting, orient the internal guide tube toward the substrate surface, and gently dispense around 30 mL of water per pot into the system. After watering, promptly reconnect the air-pump tubing to the air filter to resume continuous sterile airflow.

Note: This procedure can be performed outside a sterile workspace, provided that both filters remain securely attached to their respective GnotoBag ports and the connections downstream of the filters remain closed. During irrigation, the air filter remains attached to the gas-exchange port, maintaining a filtration barrier while the air-pump tubing is disconnected. Reconnect the air-pump tubing promptly after watering to restore continuous airflow and positive pressure.

14. Regularly rotate GnotoBag positions to minimize differential plant growth due to positional effects.
15. For axenic conditions, monitor the R2A and PDA agar-containing beakers placed inside each GnotoBag unit throughout plant growth for visible signs of contamination. At the final experimental harvest, perform an additional sterility check alongside plant sampling. To do so, open the GnotoBags in a pre-cleaned laminar flow hood or biosafety cabinet. Transfer 1 mL of standing liquid from the bottom of each Microbox onto R2A and PDA agar plates. Using a sterile inoculation loop, collect a small amount of substrate and spread it evenly onto separate R2A and PDA agar plates. Incubate all plates at room temperature for at least 1 week and monitor for microbial contamination. Plant samples may be processed while the sterility tests are pending. If contamination is detected, exclude the corresponding plant samples and any resulting data from the final analysis.

## TROUBLESHOOTING

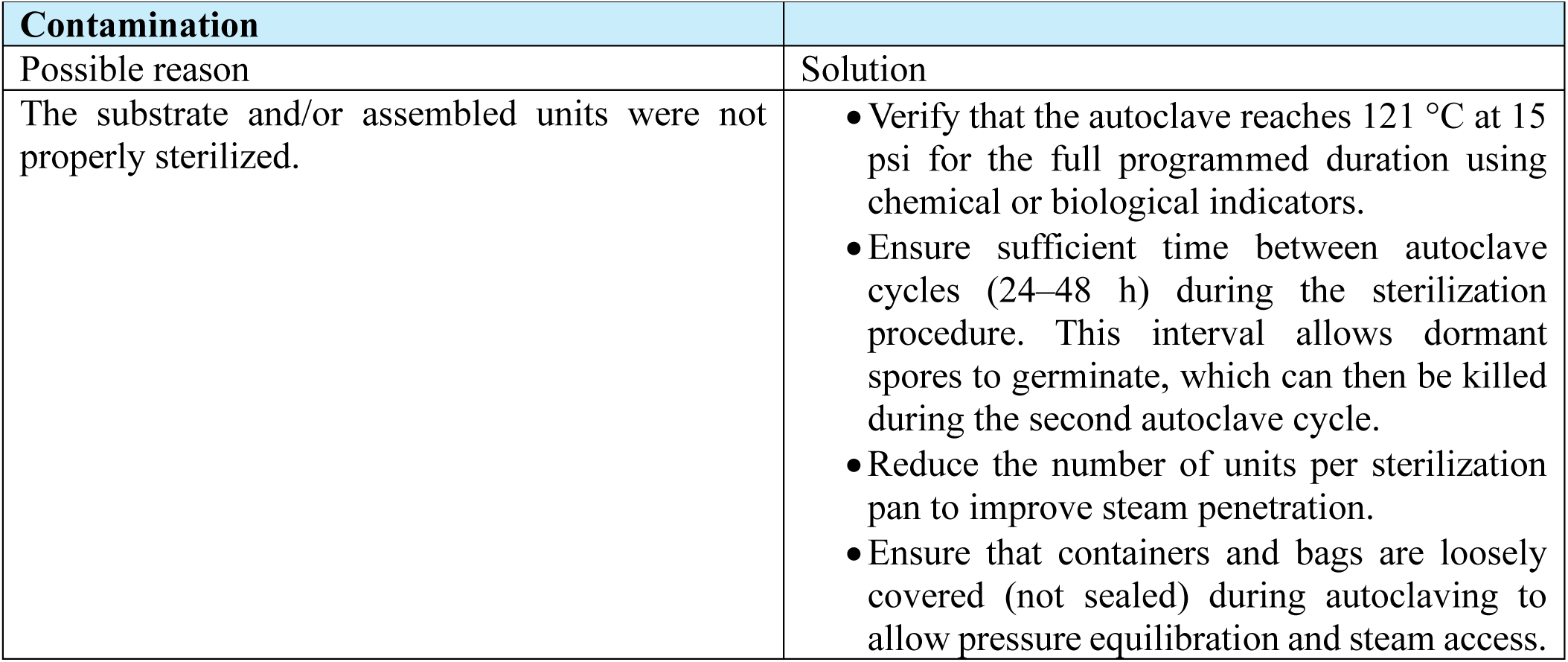
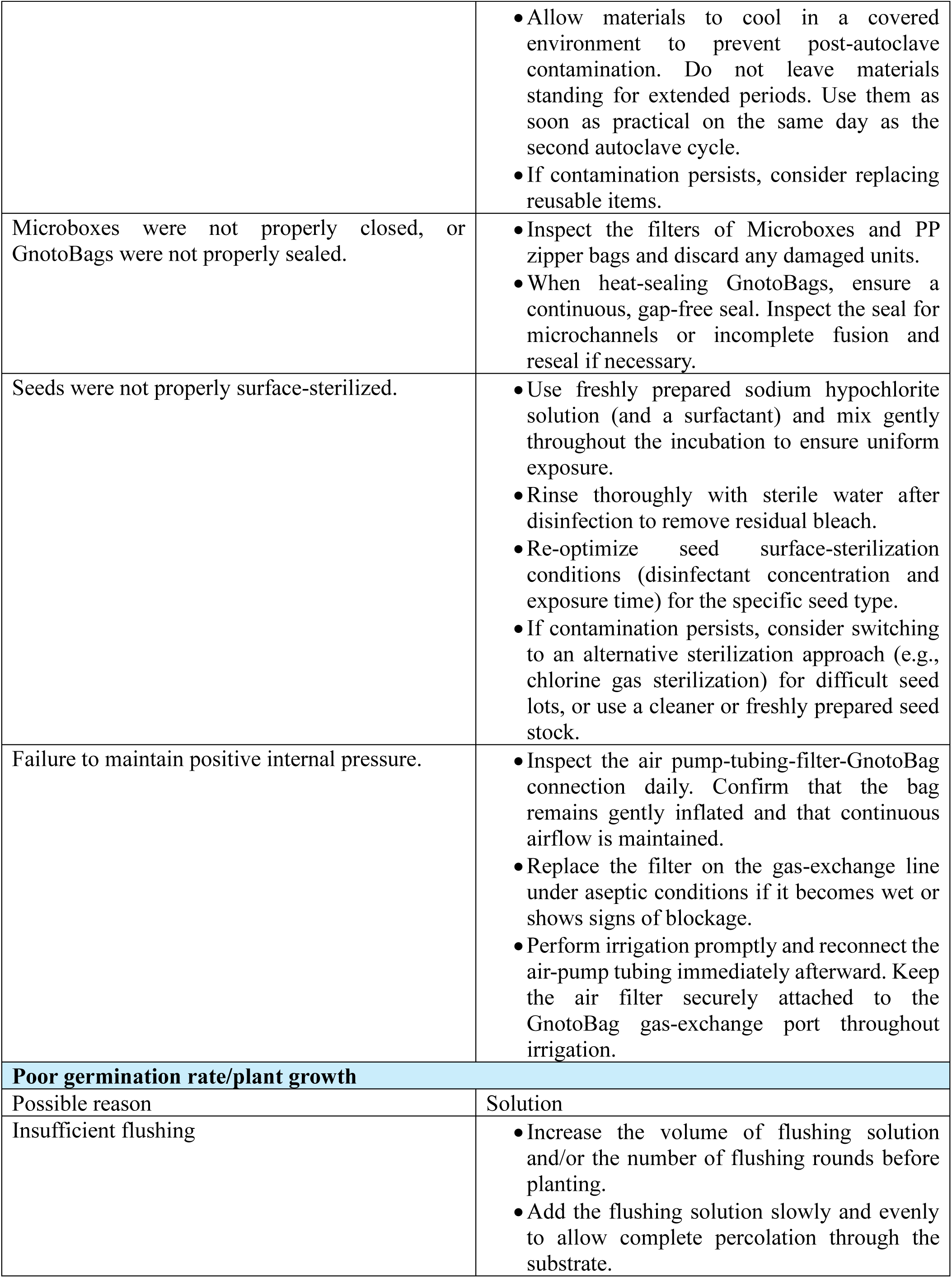
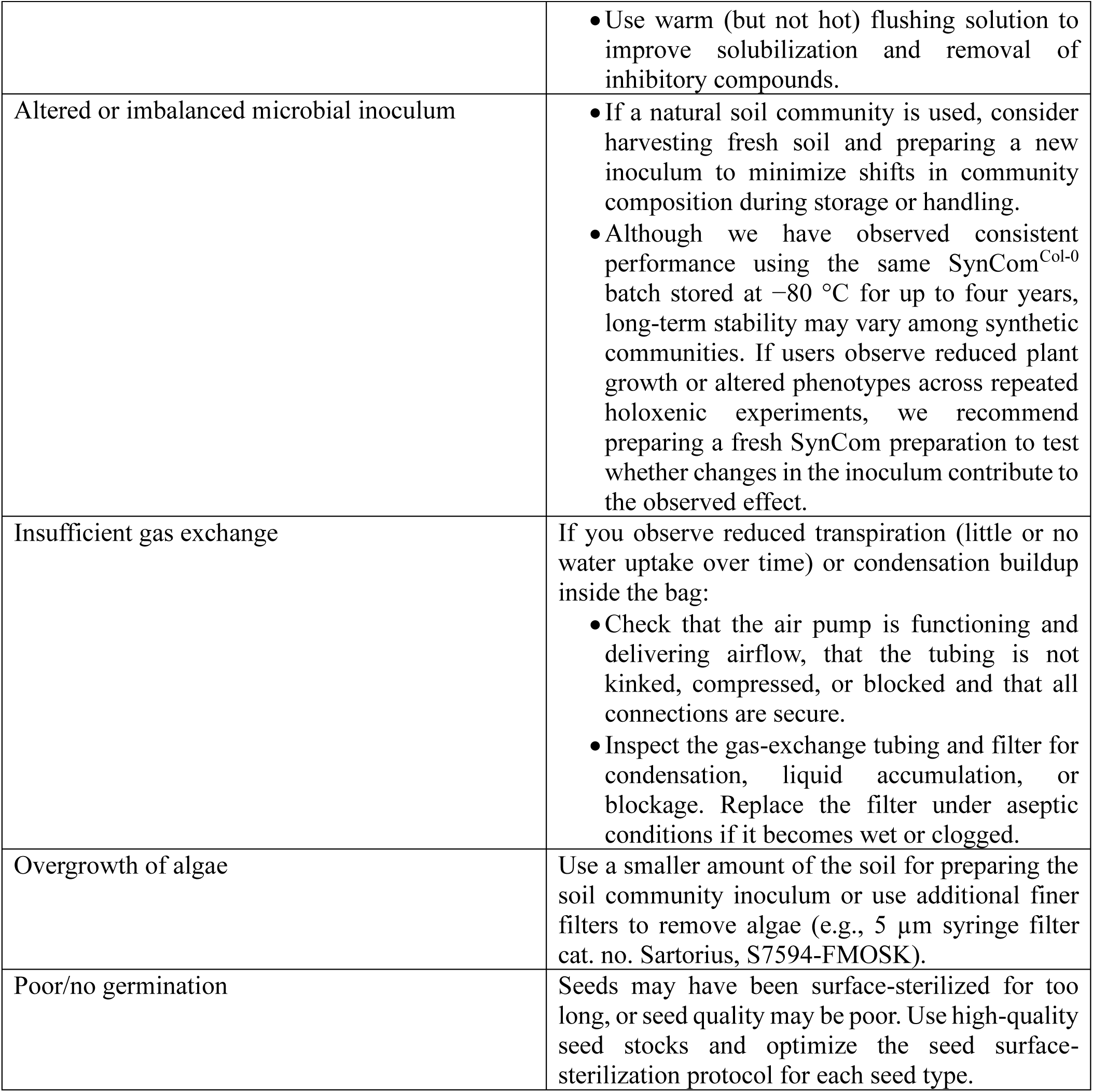

| **Contamination** |  |
|---|---|
| Possible reason | Solution |
| The substrate and/or assembled units were not properly sterilized. | • Verify that the autoclave reaches 121 °C at 15 psi for the full programmed duration using chemical or biological indicators.<br>• Ensure sufficient time between autoclave cycles (24–48 h) during the sterilization procedure. This interval allows dormant spores to germinate, which can then be killed during the second autoclave cycle.<br>• Reduce the number of units per sterilization pan to improve steam penetration.<br>• Ensure that containers and bags are loosely covered (not sealed) during autoclaving to allow pressure equilibration and steam access.<br>• Allow materials to cool in a covered environment to prevent post-autoclave contamination. Do not leave materials standing for extended periods. Use them as soon as practical on the same day as the second autoclave cycle.<br>• If contamination persists, consider replacing reusable items. |
| Microboxes were not properly closed, or GnotoBags were not properly sealed. | • Inspect the filters of Microboxes and PP zipper bags and discard any damaged units.<br>• When heat-sealing GnotoBags, ensure a continuous, gap-free seal. Inspect the seal for microchannels or incomplete fusion and reseal if necessary. |
| Seeds were not properly surface-sterilized. | • Use freshly prepared sodium hypochlorite solution (and a surfactant) and mix gently throughout the incubation to ensure uniform exposure.<br>• Rinse thoroughly with sterile water after disinfection to remove residual bleach.<br>• Re-optimize seed surface-sterilization conditions (disinfectant concentration and exposure time) for the specific seed type.<br>• If contamination persists, consider switching to an alternative sterilization approach (e.g., chlorine gas sterilization) for difficult seed lots, or use a cleaner or freshly prepared seed stock. |
| Failure to maintain positive internal pressure. | • Inspect the air pump-tubing-filter-GnotoBag connection daily. Confirm that the bag remains gently inflated and that continuous airflow is maintained.<br>• Replace the filter on the gas-exchange line under aseptic conditions if it becomes wet or shows signs of blockage.<br>• Perform irrigation promptly and reconnect the air-pump tubing immediately afterward. Keep the air filter securely attached to the GnotoBag gas-exchange port throughout irrigation. |
| **Poor germination rate/plant growth** |  |
| Possible reason | Solution |
| Insufficient flushing | • Increase the volume of flushing solution and/or the number of flushing rounds before planting.<br>• Add the flushing solution slowly and evenly to allow complete percolation through the substrate.<br>• Use warm (but not hot) flushing solution to improve solubilization and removal of inhibitory compounds. |
| Altered or imbalanced microbial inoculum | • If a natural soil community is used, consider harvesting fresh soil and preparing a new inoculum to minimize shifts in community composition during storage or handling.<br>• Although we have observed consistent performance using the same SynCom<sup>Col-0</sup> batch stored at −80 °C for up to four years, long-term stability may vary among synthetic communities. If users observe reduced plant growth or altered phenotypes across repeated holoxenic experiments, we recommend preparing a fresh SynCom preparation to test whether changes in the inoculum contribute to the observed effect. |
| Insufficient gas exchange | If you observe reduced transpiration (little or no water uptake over time) or condensation buildup inside the bag:<br>• Check that the air pump is functioning and delivering airflow, that the tubing is not kinked, compressed, or blocked and that all connections are secure.<br>• Inspect the gas-exchange tubing and filter for condensation, liquid accumulation, or blockage. Replace the filter under aseptic conditions if it becomes wet or clogged. |
| Overgrowth of algae | Use a smaller amount of the soil for preparing the soil community inoculum or use additional finer filters to remove algae (e.g., 5 µm syringe filter cat. no. Sartorius, S7594-FMOSK). |
| Poor/no germination | Seeds may have been surface-sterilized for too long, or seed quality may be poor. Use high-quality seed stocks and optimize the seed surface-sterilization protocol for each seed type. |

## ANTICIPATED RESULTS

In our previous work, the FlowPot and GnotoPot systems enabled controlled studies of plant-microbiota interactions in peat-based growth environments that more closely resemble conventional substrate-based cultivation than agar- or liquid-based systems [8]. These systems have supported a range of plant-microbiome studies [15, 16]. They also provided a starting point for further optimization. In particular, several practical needs motivated the development of the GnotoBag workflow described here: adapting the substrate to support seed germination and growth of additional plant species, increasing aerial space to accommodate vertical shoot growth, and extending gnotobiotic cultivation to later developmental stages, including seed-to-seed growth in *Arabidopsis*.

The procedures described in this protocol establish an integrated GnotoBag workflow that combines substrate adaptation and preparation with an expanded, aerated bag-based growth unit (Figs. 1 and 2). Together, these modifications provide a flexible platform for investigating plant-microbiome interactions across different plant species and developmental stages under controlled microbial conditions. After substrate adaptation, the loose peat-vermiculite mixture should remain evenly packed in the 2-inch polypropylene pots after sterilization, flushing, and initial irrigation. Compared with the fixed Jiffy-7 pellet format used in the original GnotoPot workflow, this pot-based substrate format allows the substrate volume, pot size, and seed placement to be adjusted more easily for different plant species. Under the conditions described here, the substrate-filled pots support reproducible seed germination and early plant growth under both axenic and holoxenic conditions (Fig. 3a). These prepared pots can be maintained within Microboxes for shorter-term assays or incorporated into GnotoBag units for extended cultivation with additional headspace, aeration, and sterile water or nutrient delivery (Fig. 4a).

To demonstrate these applications, we performed Procedure 1 and Procedure 2 to generate axenic and holoxenic plants under controlled microbial conditions. Procedure 1 was used to generate *Arabidopsis* plants in substrate-filled pots maintained within Microboxes for short-term growth and immune-response assays (Fig. 3). Procedure 2 was used to generate *Arabidopsis*, tomato, and rice plants in the GnotoBag system for extended growth and crop cultivation assays (Figs. 4–6). Holoxenic plants were inoculated with SynCom^Col-0^ or soil slurry, depending on the experiment, whereas axenic controls were treated with heat-killed SynCom^Col-0^ or heat-killed soil slurry. SynCom^Col-0^ [14] was generated as described in Box 1. The soil slurry was prepared from soil collected from an agricultural field at Michigan State University, East Lansing, MI, USA. For the GnotoBag experiments, samples for endpoint sterility checks were collected at the final experimental harvest. Only axenic units with no microbial growth detected after at least 1 week of incubation were included in the analyses.

In substrate-filled pots maintained within Microboxes, *Arabidopsis* seeds are expected to germinate and grow reproducibly under the recommended growth chamber conditions. In representative experiments, axenic and holoxenic *Arabidopsis* plants were grown for 2 and 4 weeks, and axenic plants remained free of detectable contamination when tested using the sterility verification procedure described in this protocol. Representative images of 2- and 4-week-old *Arabidopsis* rosettes are shown in Fig. 3a. These results show that replacing the fixed Jiffy-7 pellet format described in the original GnotoPot system with a loose peat-vermiculite substrate did not compromise *Arabidopsis* growth under controlled microbial conditions. Plant samples generated from this workflow can be used for phenotyping assays, microbial community profiling, and host-response analyses to evaluate the effects of microbiota on plant growth, immunity, and microbiome assembly.

Previous work from our lab using the GnotoPot system showed that axenic and holoxenic *Arabidopsis* plants differ in immune-related phenotypes [9]. To determine whether the substrate-filled pot workflow described here preserves this application, we performed flg22-induced reactive oxygen species (ROS) burst assays using 4-week-old *Arabidopsis* plants grown under axenic and holoxenic conditions. Consistent with previous observations in the GnotoPot system, axenic *Arabidopsis* plants showed lower flg22-induced ROS production during the 1 h assay period and lower total ROS accumulation than holoxenic plants (Fig. 3b). These representative results indicate that substrate-filled pots maintained within Microboxes support reproducible *Arabidopsis* growth and that plants grown in this system exhibit microbiota-dependent immune-response phenotypes similar to those observed in plants grown using the GnotoPot system.

The GnotoBag system is expected to provide additional headspace and continuous sterile aeration for long-term plant growth. A properly assembled GnotoBag should form a sealed, inflated unit equipped with two sterile 0.22 µm-filtered luer ports: one for continuous aeration and the other for sterile water or nutrient supplementation. Under normal operation, the GnotoBag typically inflates within approximately 10 min and remains continuously aerated throughout plant growth. The tubing, luer connections and filters should remain securely connected and free of leaks or kinks during cultivation.

In representative *Arabidopsis* experiments, plants grown in the GnotoBag system remained healthy under both axenic and holoxenic conditions (Fig. 4a). In addition to supporting vegetative growth comparable to that observed in FlowPot and GnotoPot systems, the GnotoBag system allowed *Arabidopsis* plants to progress to flowering and seed production under controlled microbial conditions (Fig. 4a). To characterize the environment inside the GnotoBag, we monitored air temperature and relative humidity using a wireless sensor placed inside the bag. Relative humidity decreased from approximately 100% at the start of aeration to approximately 60% within 6 d, while air temperature followed the programmed daily temperature cycle of the growth chamber, ranging from approximately 18 to 23 °C throughout the 14-d monitoring period. The growth chamber was programmed to 18 °C at night and 20 °C during the day to compensate for additional heat generated by the lighting system (Fig. 4b). These measurements characterize the internal conditions of the monitored GnotoBag but do not establish equivalence with the surrounding growth environment. These observations highlight the importance of monitoring the internal GnotoBag environment and adjusting growth chamber settings as needed to meet the requirements of the plant species and developmental stage.

The expanded headspace of the GnotoBag system also enables gnotobiotic cultivation of larger plant species. Tomato plants were grown under axenic and holoxenic conditions in the GnotoBag system following the procedures described in this protocol. Representative images were taken 4 weeks after sowing (Fig. 5a). We also performed flg22-induced ROS burst assays using axenic and holoxenic tomato leaves. Similar to the pattern observed in *Arabidopsis*, axenic tomato plants showed lower ROS production and lower total ROS accumulation than holoxenic tomato plants (Fig. 5b). These results demonstrate that GnotoBag can support tomato growth under controlled microbial conditions and can be used to assess microbiota-associated immune responses in this crop species.

Rice was used as an additional crop species to evaluate the broader applicability of the GnotoBag system. In this study, seeds of the rice cultivar Kitaake were surface-sterilized as described in this protocol and sown into the peat-based substrate. Plants were maintained in GnotoBags under growth chamber or greenhouse conditions, with sterile water and nutrient supplementation provided through the irrigation line as needed. Representative images of 4-week-old rice plants grown in the GnotoBag system are shown in Fig. 6a.

To assess host immune responses in rice, leaves were collected for RNA extraction and reverse transcription quantitative PCR (RT-qPCR) analysis. We quantified the relative transcript levels of *OsFLS2* and *OsPR1b*, two rice defense-associated marker genes. *OsFLS2* encodes a rice flagellin receptor involved in flg22 perception [17], whereas *OsPR1b* belongs to the rice PR1 gene family and has been characterized in relation to defense-associated signaling and stress responses [18, 19]. Holoxenic rice plants grown in the GnotoBag system showed higher relative transcript levels of both genes than axenic rice plants (Fig. 6b), demonstrating microbiota-dependent differences in defense-associated gene expression. Similar differences in defense-associated gene expression between axenic and holoxenic conditions have previously been reported in *Arabidopsis* [9]. These representative results indicate that GnotoBag can support gnotobiotic cultivation of rice and can be used to examine microbiota-associated host responses in crop plants. Collectively, these examples demonstrate the versatility of the GnotoBag system for maintaining plants under controlled microbial conditions across different plant species and developmental stages. Users can expect healthy plant growth together with sufficient space and accessibility for sterile irrigation, nutrient supplementation, and downstream phenotyping. Importantly, the successful cultivation of crop species such as tomato and rice extends the utility of gnotobiotic systems beyond *Arabidopsis* and enables plant-microbiota interactions to be investigated in agriculturally relevant hosts and over longer developmental periods. This expanded capacity may facilitate studies of microbiome effects on crop growth, nutrient use, and yield-related traits, thereby helping to bridge mechanistic gnotobiotic research with crop improvement and broader agricultural applications.

## ACKNOWLEDGEMENTS

Figures 1 and 2 were partially created with BioRender.com. Some schematic elements, including the Nalgene polypropylene sterilizing pan covered with sterilization wrap in Fig. 1 and the schematic elements in Fig. 2b, c, were drafted with assistance from ChatGPT and subsequently reviewed, edited, and finalized by the authors for accuracy. No experimental data were generated, analyzed, or modified using AI tools. This work was supported by the Duke University Science and Technology Initiative to S.Y.H. S.Y.H. is an Investigator of the Howard Hughes Medical Institute. N.W. is supported by a fellowship from the Institute of Plant Protection, Chinese Academy of Agricultural Sciences, and by the China Postdoctoral Science Foundation (2022M723452).

## AUTHOR CONTRIBUTIONS

N.W., Y.T.C. and B.C.P. collaborated on the conception, design and/or optimization of various steps of the GnotoBag system with input from S.Y.H. N.W., Y.T.C. and S.Y.H. wrote and finalized the manuscript with input from all coauthors.

